# Vertical transmission at the pathogen-symbiont interface: *Serratia symbiotica* and aphids

**DOI:** 10.1101/2020.09.01.279018

**Authors:** Julie Perreau, Devki J. Patel, Hanna Anderson, Gerald P. Maeda, Katherine M. Elston, Jeffrey E. Barrick, Nancy A. Moran

## Abstract

Many insects possess beneficial bacterial symbionts that occupy specialized host cells and are maternally transmitted. As a consequence of their host-restricted lifestyle, these symbionts often possess reduced genomes and cannot be cultured outside hosts, limiting their study. The bacterial species *Serratia symbiotica* was originally described by noncultured strains that live as mutualistic symbionts of aphids and are vertically transmitted through transovarial endocytosis within the mother’s body. More recently, culturable strains of *S. symbiotica* were discovered that retain a larger set of ancestral *Serratia* genes, are gut pathogens in aphid hosts, and are principally transmitted via a fecal-oral route. We find that these culturable strains, when injected into pea aphids, replicate in the hemolymph and are pathogenic. Unexpectedly, they are also capable of maternal transmission via transovarial endocytosis: using GFP-tagged strains, we observe that pathogenic *S. symbiotica*, but not *Escherichia coli*, are endocytosed into early embryos. Furthermore, pathogenic *S. symbiotica* strains are compartmentalized into specialized aphid cells in a similar fashion to mutualistic *S. symbiotica* strains during later stages of embryonic development. Thus, cultured, pathogenic strains of *S. symbiotica* have the latent capacity to transition to lifestyles as mutualistic symbionts of aphid hosts. This capacity is blocked by pathogenicity: their hosts die before infected progeny are born. To transition into stably inherited symbionts, culturable *S. symbiotica* strains may need to adapt to regulate their titer, limit their pathogenicity, and/or provide benefits to aphids that outweigh their cost.

**Importance:** Insects have evolved various mechanisms to reliably transmit their beneficial bacterial symbionts to the next generation. Sap-sucking insects, including aphids, transmit symbionts by endocytosis of the symbiont into cells of the early embryo within the mother’s body. Experimental studies of this process are hampered by the inability to culture or genetically manipulate host-restricted, symbiotic bacteria. *Serratia symbiotica* is a bacterial species that includes strains ranging from obligate, heritable symbionts to culturable gut pathogens. We demonstrate that culturable *S. symbiotica* strains, that are aphid gut pathogens, can be maternally transmitted by endocytosis. Cultured *S. symbiotica* therefore possess a latent capacity for evolving a host-restricted lifestyle and can be used to understand the transition from pathogenicity to beneficial symbiosis.

## Introduction

Host-associated bacteria can be placed on a continuum ranging from parasitic to mutualistic. Mutualistic symbionts often arise from pathogenic ancestors and rarely revert back to pathogenicity (1). This one-way transition is expected when vertical transmission, usually from mother to offspring, replaces horizontal transmission as the dominant route of new infection, causing symbionts to benefit from host reproduction and thereby to face strong selection for avirulence (2–4). In insects, the reliable vertical transmission of mutualistic symbionts can be accomplished by mechanisms that are external, including the placement of symbionts or symbiont-containing capsules on the egg surface, or internal, via transovarial transmission (5, 6).

Transovarial transmission, the transfer of maternal symbionts to eggs or embryos within the mother’s body, is common in obligate symbioses where symbionts occupy special host cells (bacteriocytes) within the body cavity (7). Individual bacteria may be transferred, as in aphids, cicadas, leafhoppers, cockroaches, and bedbugs, or entire bacteriocytes may be transferred, as in whiteflies (7–10). In ancient symbioses with exclusively maternal transmission, hosts appear to control transmission, as the symbionts involved often possess reduced genomes devoid of pathogenicity factors that would allow them to invade host cells (11). However, host mechanisms for transmission may depend on bacterial factors for inter-partner recognition, and thereby limit which bacterial species or strains can make the transition from pathogenicity to commensal or mutualistic symbioses.

Aphids (Hemiptera: Sternorrhyncha: Aphidoidea) are a clade of roughly 5,000 species that feed exclusively on nutrient-poor plant sap and depend on the primary bacterial symbiont *Buchnera aphidicola* for biosynthesis of essential amino acids missing from their diet (12–16). *B. aphidicola* have been transovarially transmitted in aphids for over 160 million years (17). They occupy bacteriocytes, possess highly reduced genomes, and cannot persist outside of their hosts (11, 18). In addition to *B. aphidicola*, many aphids also host secondary symbionts, such as *Serratia symbiotica, Candidatus* Hamiltonella defensa, *Candidatus* Regiella insecticola, and *Candidatus* Fukatsuia symbiotica (19–22). Due to their more recent associations with aphids, secondary symbionts provide an opportunity to understand the transition to host-restricted lifestyles.

*S. symbiotica* strains have evolved diverse associations with aphids. They range from pathogens and facultative mutualists to obligate mutualists that co-reside with *B. aphidicola* (23–32). The genome sizes and gene contents of *S. symbiotica* strains reflect this variation in lifestyle, ranging from larger genomes similar to those of free-living *Serratia* (31, 33), to highly reduced genomes similar to those of *B. aphidicola* and other obligate symbionts (24, 26, 27, 32). The first descriptions of *S. symbiotica*, initially named Pea Aphid Secondary Symbiont (PASS) or the “R-type” symbiont, were of strains that occupied secondary bacteriocytes, sheath cells, and hemolymph (34, 19). Unlike *B. aphidicola*, these *S. symbiotica* strains are not required by their hosts; they provide secondary benefits, such as protection against heat stress (23, 35), and against parasitoid wasps (36). However, similar to *B. aphidicola*, they are host-restricted and vertically transmitted by a transovarial route (8, 37, 38). A detailed study showed that a mutualistic *S. symbiotica* strain, present in the hemolymph, migrates to early embryos and is endocytosed with *B. aphidicola* into the syncytium, i.e., a specialized, multinucleated cell of the early embryo (8). In the pea aphid, *B. aphidicola* and *S. symbiotica* are later segregated into distinct bacteriocytes.

Recently, strains of pathogenic *S. symbiotica* have been discovered living in the guts of *Aphis* species collected in Belgium and Tunisia (25, 29–31). These strains are hypothesized to resemble ancestors of facultative and co-obligate *S. symbiotica* strains (39, 40). In contrast to previously studied *S. symbiotica* strains, their primary transmission route appears to be horizontal, through honeydew (feces) and host plant phloem (39, 40). Also, in contrast to previously studied strains, these *S. symbiotica* can be cultured axenically. *S. symbiotica* CWBI-2.3^T^, from the black bean aphid (*Aphis fabae*) (25), retains a larger gene set and larger genome (3.58 Mb) (33) than do facultative *S. symbiotica* strains Tucson and IS (2.78 and 2.82 Mb, respectively) (41, 42).

In this work, we investigate whether cultured *S. symbiotica* strains are capable of vertical transmission similar to facultative or obligate symbionts. We isolated a new *S. symbiotica* strain, HB1, that shares many features with CWBI-2.3^T^ but is notably less pathogenic. We examined the capacity of each of these strains to colonize hemolymph of the pea aphid (*Acyrthosiphon pisum*) and access embryos through the transovarial route described for *B. aphidicola* (8). Both strains are endocytosed into embryos, but pathogenic strains kill their hosts before the birth of offspring infected by the transovarial route. Using GFP-tagged strains, we addressed whether transovarial transmission is open to any bacterial cell that comes into contact with the embryo, or whether this process involves specific partner recognition. We find that *Escherichia coli* cannot colonize embryos, despite achieving a high titer within hosts. Thus, the endocytosis step required for transovarial transmission limits the taxonomic range of bacteria that can readily evolve to become aphid symbionts.

## Results

### Pathogenic *S. symbiotica* form a distinct group closely related to mutualistic strains

We examined the evolutionary relationships of all *S. symbiotica* with publicly available complete genome sequences. To this list, we added a recently isolated and sequenced strain, designated *S. symbiotica* HB1, from the melon aphid (*Aphis gossypii*). Our phylogenetic analysis was based on 176 shared orthologous genes and was rooted with outgroups that included other *Serratia* and more distant *Enterobacterales* species (Figure 1, Figure S1, Table S1). *S. symbiotica* is split into two clades, as previously reported (43). Clade A is comprised of strains that act as pathogens, mutualists, or co-obligate symbionts in aphids from across the family Aphididae, while clade B is composed of strains that live only as co-obligate symbionts in aphids of the subfamily Lachninae. Clade A strains possess a range of genome sizes (1.54–3.58 Mb), GC content (47.9–52.5%), host species, and lifestyles. Within clade A, the cultured, gut pathogen strains (CWBI-2.3^T^, HB1, Apa8A1, and 24.1) form a clade closely related to vertically transmitted mutualist strains (Tucson, IS, MCAR-56S, AURT-53S). Although average nucleotide identities are high (96.0–98.6%) across these strains (Table S2), the gut pathogen strains retain larger genomes (3.09–3.58 Mb) and more *Serratia*-specific marker genes than non-pathogenic, maternally transmitted *S. symbiotica* (0.65–2.82 Mb) (Figure 1, Table S1). Together, these observations suggest that both lifestyles have recently emerged from a common, presumably pathogenic ancestor.

**Figure 1.**
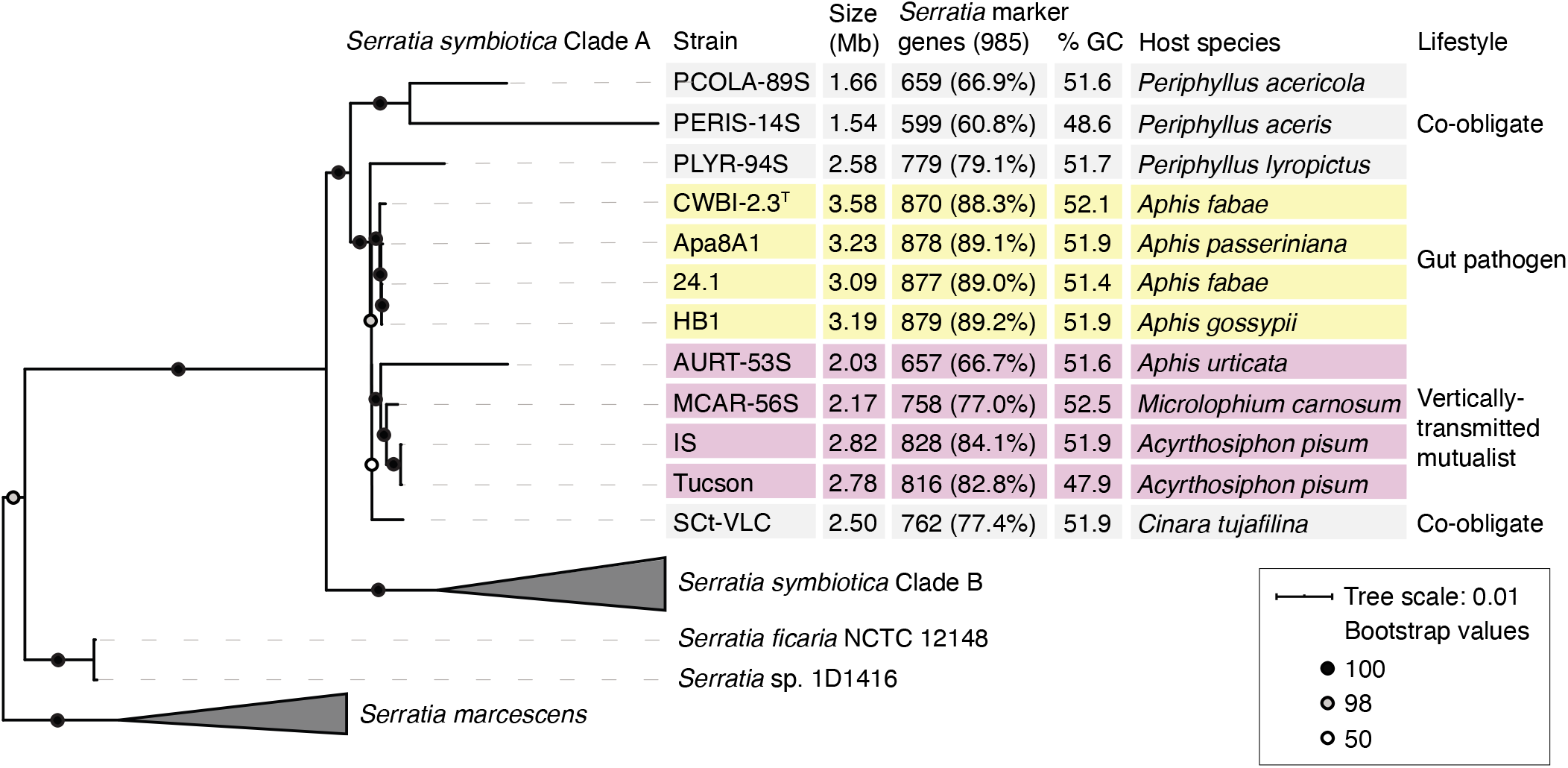
Phylogeny and gene content evolution of *S. symbiotica*. (A) Maximum likelihood phylogeny of *S. symbiotica*, based on concatenation of 176 single-copy orthologs (56,881 amino acid positions) shared across *S. symbiotica*, other *Serratia* species, *E. coli, Yersinia pestis*, and *Salmonella enterica*. Genomes used for the phylogeny and relevant references for *S. symbiotica* genome features are listed in Table S1. Bootstrap values are indicated with symbols on nodes. The complete phylogeny is presented in Figure S1.

### Cultured *S. symbiotica* strains are pathogenic when injected into pea aphid hemolymph

Based on previous studies, *S. symbiotica* CWBI-2.3^T^ acts as a gut pathogen in its original host, the black bean aphid (40), but appears to be less pathogenic in the gut of pea aphids (44). In both hosts, *S. symbiotica* CWBI-2.3^T^ appears to be restricted to the gut and is not observed infecting hemolymph. To determine whether *S. symbiotica* CWBI-2.3^T^ and HB1 can persist and act as pathogens in the hemolymph of pea aphids, and to determine if these strains are capable of vertical transmission to offspring, we injected fourth instar pea aphids with tagged strains CWBI-2.3^T^-GFP or HB1-GFP. Then, we tracked aphid survival, vertical transmission rates, and bacterial titer over time. For comparison, we simultaneously performed injections of *S. marcescens* Db11, a known insect pathogen (45), injections of hemolymph from pea aphids infected with facultative, vertically transmitted *S. symbiotica* Tucson (41), and injections of buffer as a negative control. We found that *S. symbiotica* CWBI-2.3^T^-GFP and HB1-GFP act as pathogens in pea aphid hemolymph. The survival rates of aphids injected with *S. symbiotica* CWBI-2.3^T^-GFP, *S. symbiotica* HB1-GFP, or *S. marcescens* Db11 were much lower than those of aphids injected with buffer or with *S. symbiotica* Tucson (P < 0.001, Cox Proportional Hazards Model) (Figure 2A).

**Figure 2.**
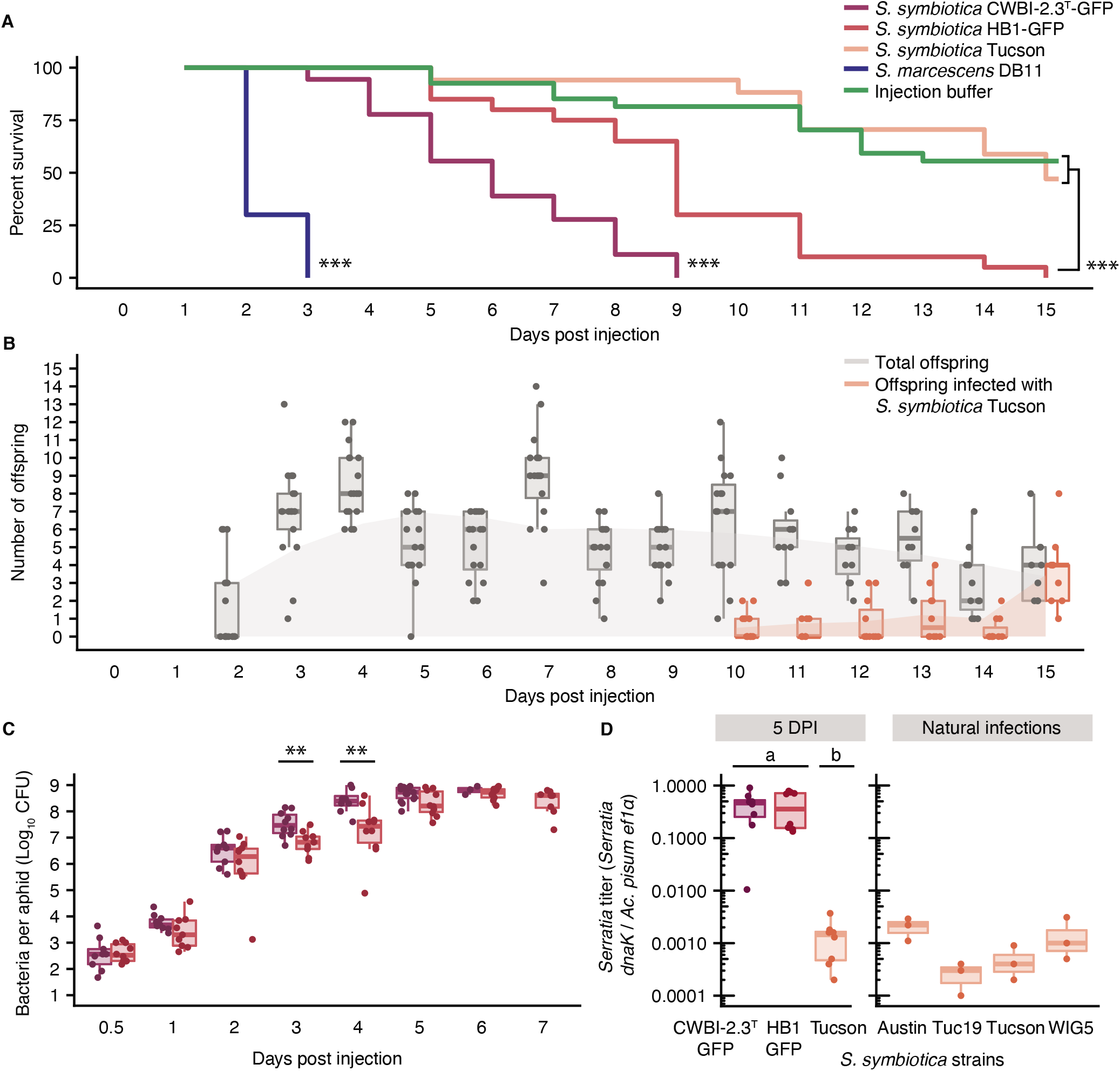
Infection dynamics of *S. symbiotica* strains CWBI-2.3^T^ and HB1 injected into hemolymph of fourth instar pea aphids. (A) Survival curves of pea aphids injected with cultured *S. marcescens* Db11, recombinant *S. symbiotica* CWBI-2.3^T^-GFP, or recombinant *S. symbiotica* HB1-GFP, with hemolymph from *S. symbiotica* Tucson, or with injection buffer. *** P < 0.001, Cox Proportional Hazards Model. (B) Fecundity and transmission of *S. symbiotica* for adults injected with *S. symbiotica* Tucson. Boxplots in gray depict the total offspring, including infected and uninfected offspring, per adult. Boxplots in orange depict total offspring infected with *S. symbiotica* Tucson per adult. Filled curves represent a LOESS regression and display the increase in offspring infected with *S. symbiotica* Tucson from 10 DPI to 15 DPI. (C) Bacterial titer of recombinant *S. symbiotica* strains CWBI-2.3^T^-GFP and HB1-GFP, obtained by spot-plating dilutions from whole aphids. ** p < 0.01, Wilcoxon rank-sum test. (D) Relative titer of *S. symbiotica* across treatment groups at 5 DPI (left) and for 7-day-old pea aphids from lab-reared clonal lines that are naturally infected with vertically transmitted *S. symbiotica* strains, established from collections in Austin in 2014 (Austin), Tucson in 2019 (Tuc19), Tucson in 1999 (Tucson), and Wisconsin in 2017 (WIG5) (right). Relative titer was calculated as the copy number of *S. symbiotica dnaK* normalized by copy number of the single copy *Acyrthosiphon pisum* gene *ef1a*. CWBI-2.3^T^ and HB1 samples used for qPCR are the same as those used to determine bacterial titer by spot-plating. Letters a and b denote two groups of strains that have significantly different titers from one another and indistinguishable titers within groups at α = 0.01 by the Kruskal-Wallis test followed by Dunn’s multiple comparison test.

Typically, *S. symbiotica* CWBI-2.3^T^ infects by a fecal-oral route or from plants (40). To determine if *S. symbiotica* CWBI-2.3^T^-GFP and HB1-GFP can be transmitted to offspring via the transovarial route, as for mutualistic symbiotic strains, we collected newborn offspring of surviving aphids every 24 hours and screened them for *S. symbiotica* by plating aphids and noting presence or absence of colony-forming units (CFU). As transovarial transmission of symbionts occurs early in embryonic development, there is a delay between injection and the birth of infected offspring (34). To determine when after injection we should begin to observe offspring infected by transovarial transmission, we injected mutualistic *S. symbiotica* Tucson and monitored its transmission by sampling offspring and using PCR to screen for the presence of *S. symbiotica*. Transmission of *S. symbiotica* Tucson was identified in newborn offspring starting at 10 days post-injection (DPI) (Figure 2B). The proportion of infected offspring per mother increased over time, reaching 100% for most mothers by 15 DPI (Figure 2B). In contrast to aphids injected with *S. symbiotica* Tucson, aphids injected with *S. symbiotica* CWBI-2.3^T^ did not survive past 9 DPI (Figure 2A), and aphids injected with *S. symbiotica* HB1 showed reduced fecundity and produced only a few, non-viable offspring after 9 DPI. Therefore, successful vertical transmission of these strains appears to be limited by their pathogenicity.

We hypothesized that the virulence of *S. symbiotica* CWBI-2.3^T^-GFP and HB1-GFP was related to their titer in hemolymph. To test this hypothesis, aphids were sacrificed every 24 hours to measure the CFU per aphid. *S. symbiotica* CWBI-2.3^T^-GFP and HB1-GFP grow exponentially within aphids, plateauing at ~10^9^ CFU per aphid by 5-6 DPI (Figure 2C). The significant difference in the growth rate of CWBI-2.3^T^ and HB1 at 3 and 4 DPI may explain the difference in aphid survival rate across these treatment groups (Figure 2A). As *S. symbiotica* Tucson is not culturable, qPCR was used to compare titers of CWBI-2.3^T^-GFP and HB1-GFP to those reached by Tucson at 5 DPI, and to the titers at which other mutualist *S. symbiotica* strains persist in pea aphids reared in the lab (Figure 2D). Relative titer was calculated as the copy number of a single-copy *Serratia* gene (*dnaK*) normalized by a single-copy pea aphid gene (*ef1a*). Both *S. symbiotica* CWBI-2.3^T^-GFP and HB1-GFP reach higher titers than *S. symbiotica* Tucson at 5 DPI (Figure 2D). The titer of *S. symbiotica* Tucson at 5 DPI is comparable to the steady-state titer of *S. symbiotica* strains maintained in naturally infected, clonal pea aphids established as laboratory lines (Figure 2D).

### Cultured *S. symbiotica* strains are capable of colonizing *Ac pisum* embryos

Intergenerational transmission of *S. symbiotica* CWBI-2.3^T^ and HB1 was prevented because the pathogenicity of these strains killed mothers before any infected embryos could be born. In order to determine whether they were, nevertheless, capable of transovarial transmission, we injected aphids with *S. symbiotica* CWBI-2.3^T^, dissected ovarioles at 7 DPI, and used fluorescence *in-situ* hybridization to visualize its infection pattern relative to the primary symbiont *B. aphidicola* during embryonic development. To compare these results to the transmission pattern of a mutualist strain, we injected aphids with hemolymph from pea aphids infected with *S. symbiotica* Tucson (Figure 3A-D).

**Figure 3.**
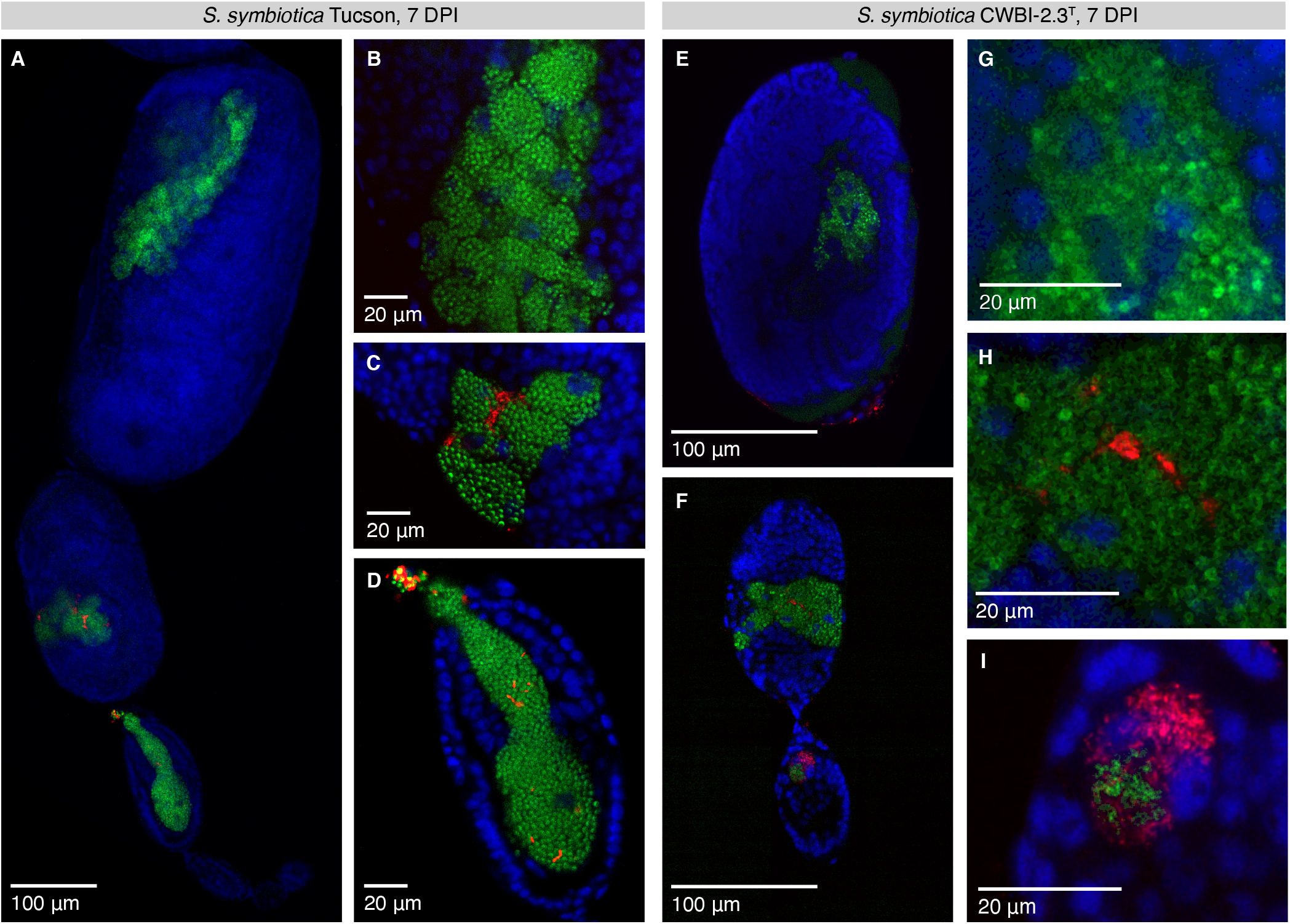
Transovarial transmission of the pathogenic strain *S. symbiotica* CWBI-2.3^T^ and the mutualistic strain *S. symbiotica* Tucson in stage 7 embryos of pea aphids. (A) *S. symbiotica* Tucson infection across embryonic stages at 7 DPI. (B) *S. symbiotica* Tucson does not infect embryos that are beyond stage 7 of development at the time of injection, as indicated by a lack of *S. symbiotica* Tucson in late-stage embryos. (C) In infected, mid-stage embryos, *S. symbiotica* Tucson cannot infect primary bacteriocytes that contain *B. aphidicola* but does infect sheath cells. (D) *S. symbiotica* Tucson enters the syncytium of an early-stage blastula with *B. aphidicola* but is underrepresented relative to *B. aphidicola* during this infection. (E-F) *S. symbiotica* CWBI-2.3^T^ infection across embryonic stages at 7 DPI. Embryos depicted in (E) and (F) derive from the same ovariole but were separated during dissection. Infection with this strain results in reduced embryonic growth, as indicated by scale bars. (G) *S. symbiotica* CWBI-2.3^T^ does not infect embryos that are beyond stage 7 of development at the time of injection, as indicated by a lack of *S. symbiotica* CWBI-2.3^T^ in late-stage embryos. (H) In infected, mid-stage embryos, *S. symbiotica* CWBI-2.3^T^ cannot infect primary bacteriocytes that contain *B. aphidicola* but does infect sheath cells. (I) *S. symbiotica* CWBI-2.3^T^ enters the syncytium of an early-stage blastula with *B. aphidicola* and is overrepresented relative to *B. aphidicola* during this infection. Blue, red, and green signal are used for host nuclei, *S. symbiotica*, and *B. aphidicola*, respectively.

Embryos injected with *S. symbiotica* Tucson display regular growth and development, reaching 400 μm in length by the time of katatrepsis (37) (Figure 3A). As previously described, *B. aphidicola* and mutualistic *S. symbiotica* are transmitted to the syncytium of stage 7 blastula from hemolymph via an endocytic process (8, 37, 46). *S. symbiotica* Tucson appears to be unable to infect embryos at later developmental stages, as embryos that were beyond stage 7 at the time of injection lack *S. symbiotica* (Figure 3B). This observation is supported by the absence of *S. symbiotica* in offspring born the first 10 days after injection (Figure 2B) (34). Those embryos exposed to *S. symbiotica* Tucson at stage 7 possess *S. symbiotica* in sheath cells, but Tucson does not invade primary bacteriocytes that contain *B. aphidicola* (Figure 3C). The endocytosis of *S. symbiotica* Tucson into stage 7 embryos occurs at the posterior end of the embryo (Figure 3D), as previously described for the closely related strain *S. symbiotica* IS (8).

Despite its pathogenicity, the infection pattern of CWBI-2.3^T^ is similar to that of the non-pathogenic Tucson strain (Figures 3E, 3F). Cells of CWBI-2.3^T^ are attached to the embryonic surface but do not infect embryos that are beyond stage 7 (Figure 3G). Following infection, CWBI-2.3^T^ is packaged similarly to the Tucson strain; both are sorted into sheath cells and cannot colonize the primary bacteriocytes that house *B. aphidicola* (Figures 3C, 3H). CWBI-2.3^T^ is endocytosed into early embryos with *B. aphidicola* (Figure 3I), as is *S. symbiotica* HB1 (Movie S1). Remarkably, at 7 DPI, CWBI-2.3^T^ greatly outnumbers *B. aphidicola* in the syncytial cell (Figure 3I). In contrast to *S. symbiotica* Tucson, infection with cultured *S. symbiotica* CWBI-2.3^T^ stunts embryonic growth, though it does not prevent progression through characteristic developmental stages (Figures 3E, 3F).

### Transovarial transmission is a specific capability of *S. symbiotica* strains

To determine if endocytosis is selective at the level of bacterial species, we tested the transmission capability of *E. coli* K-12 strain BW25113. *E. coli* is related to *B. aphidicola, S. symbiotica*, and several other mutualistic symbionts of aphids which are all within *Enterobacterales* (20). We chose strain BW25113 because it can infect the gut and hemolymph of pea aphids and kills aphids a few days post infection (47). We created the tagged *E. coli* strain BW25113-GFP and injected it into fourth instar aphids as described above. For comparison, we injected *S. symbiotica* CWBI-2.3^T^-GFP into a separate set of fourth instar aphids. *E. coli* BW25113-GFP forms a robust infection in pea aphid hemolymph, reaching comparable titers to *S. symbiotica* CWBI-2.3^T^-GFP at 5 DPI (Figure 4A). We dissected single ovarioles from ten aphids in each treatment group at 5 DPI and observed early embryos to determine infection status. Using this approach, *S. symbiotica* CWBI-2.3^T^-GFP could be seen infecting early embryos (Figures 4B, 4C, Movies S2, S3). In contrast, *E. coli* BW25113-GFP attaches to the embryonic surface, sometimes coating the entire exterior of the embryo, but was never observed endocytosing into embryos (Figures 4D, 4E, Movies S4, S5).

**Figure 4.**
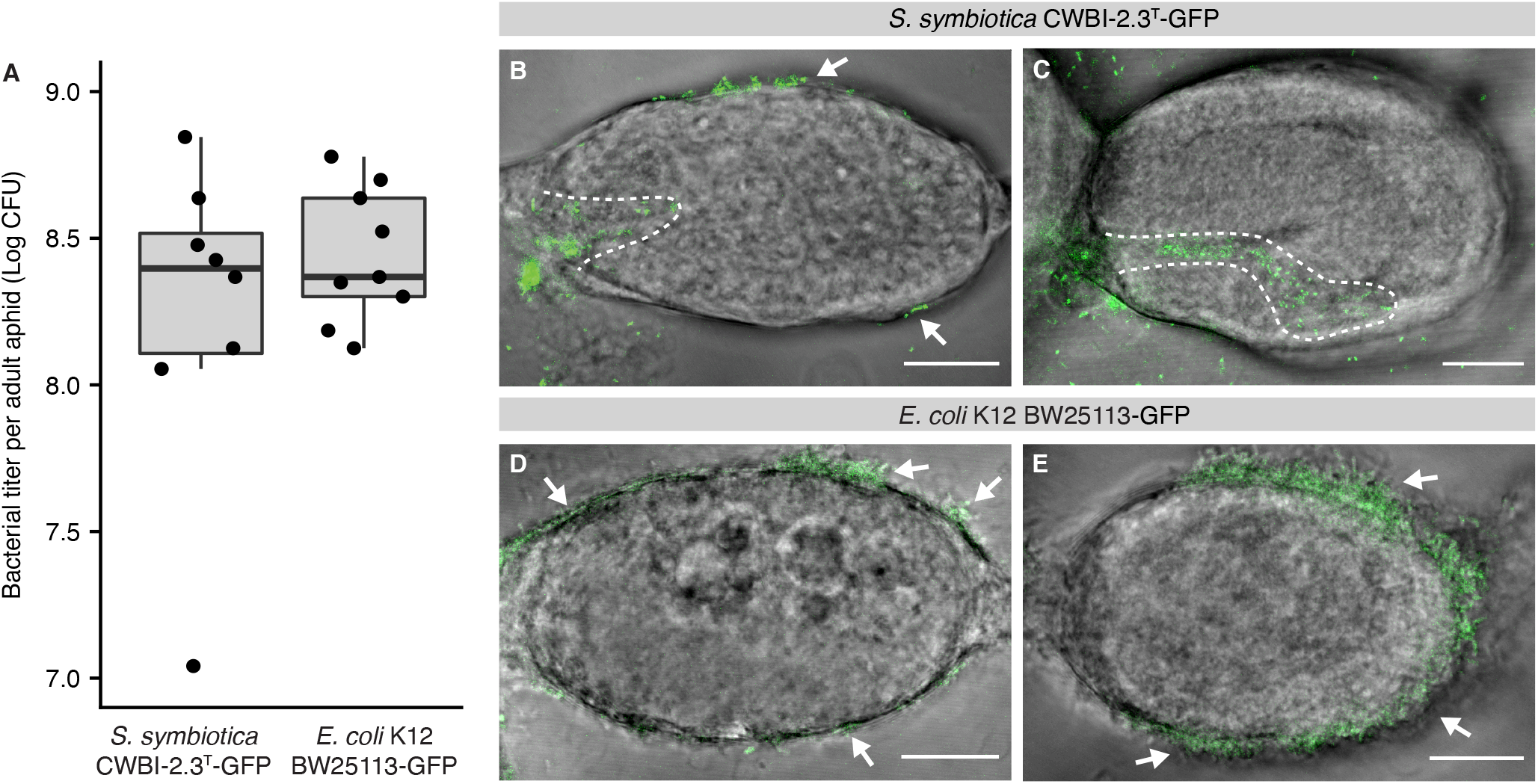
Ability of *S. symbiotica* CWBI-2.3^T^ but not *E. coli* BW25113 to achieve transovarial transmission to stage 7 embryos of pea aphids. (A) Recombinant *S. symbiotica* CWBI-2.3^T^-GFP and *E. coli* BW25113-GFP reach comparable titers 5 DPI, as determined by spot-plating. (B, C) Recombinant *S. symbiotica* CWBI-2.3^T^-GFP enters the posterior region of stage 7 blastula. Embryos were dissected and visualized at 5 DPI. Dotted lines outline *S. symbiotica* present within the embryo. (D, E) Recombinant *E. coli* BW25113-GFP does not enter into stage 7 blastula. Embryos were dissected and visualized at 5 DPI. Arrows indicate bacteria attached to the embryonic surface. Scale bars represent 20 μM.

## Discussion

Transovarial transmission is a key feature of many insect-bacterial symbioses wherein bacteria provide a benefit to their host. This transmission route is linked to irreversible bacterial transitions from pathogenicity to mutualism (2). However, many relationships that rely on transovarial transmission are ancient, and their early stages cannot be experimentally recapitulated, leaving unanswered if and how pathogenic bacteria access this transmission route. Focusing on secondary symbionts may be more useful for understanding these early transitions, as some secondary symbionts or their close relatives are culturable, genetically tractable, and can be removed from or introduced to hosts without dramatically compromising host fitness (48–50). For example, *Sodalis praecaptivus*, a close relative to *Sodalis* species found as host-restricted symbionts across diverse insects, uses quorum sensing to attenuate virulence and gain access to vertical transmission in a non-native host, the tsetse fly (51). The bacterial species *S. symbiotica* was first known as a vertically transmitted mutualist, but pathogenic strains were subsequently cultured from aphids collected in Europe and Africa (25, 29–31), and, in this study, in North America. These strains have provided a new opportunity to dissect the early steps involved in the transition to a host-restricted lifestyle. Knowing that vertical transmission is a key to this transition, we aimed to determine whether culturable, pathogenic *S. symbiotica* could access this pathway and what, if any, limitations are faced by *S. symbiotica* in this transition.

Cultured *S. symbiotica* strains are close relatives to non-culturable, vertically transmitted mutualists. However, several lines of evidence suggest these strains lack a history of maternal transmission in aphids. For one, persistent vertical transmission generally leads to the irreversible loss of genes such that they can no longer access a free-living or pathogenic lifestyle (52). Relative to vertically transmitted strains, CWBI-2.3^T^, HB1, Apa8A1, and 24.1 are culturable, maintain larger genomes, and possess more ancestral genes common to free-living *Serratia* (Figure 1). Second, these strains do not appear to undergo vertical transmission in natural infections. Following ingestion by black bean or pea aphids, *S. symbiotica* CWBI-2.3^T^ is not subsequently detected in hemolymph or embryos, but is present in the gut and in honeydew, suggesting that the dominant route of transmission for this strain is fecal-oral (29, 39, 44). We injected CWBI-2.3^T^ and HB1 into hemolymph to determine if they are capable of vertical transmission in pea aphids. Vertical transmission is theorized to be the primary force driving permanent bacterial transitions from pathogenicity to mutualism, but to do so, vertical transmission must precede mutualism (2). That pathogenic *S. symbiotica* CWBI-2.3^T^ and HB1 are endocytosed into the syncytial cell of early embryos along with *B. aphidicola* provides empirical evidence for the precedence of vertical transmission in this system. Together, these results suggest that the aphid gut has served as an access point for environmental or plant-associated *Serratia* to infect aphids and that gut pathogenicity was an ancestral lifestyle for strains that are now intracellular and mutualistic (29).

*S. symbiotica* is common to natural populations of pea aphids (34, 53), and previous 16S rRNA gene surveys have identified *S. symbiotica* in aphid tribes from across the Aphidoidea (29, 54). However, pathogenic and mutualistic strains have near-identical 16S rRNA sequences, so it is unclear how many cases represent *S. symbiotica* pathogens. To date, pathogenic strains have been cultured only from *Aphis* species. However, *S. symbiotica* CWBI-2.3^T^ can horizontally transmit across aphids feeding on the same plant (39), and can infect the guts of alternative aphid species, including the pea aphid (44), suggesting that pathogenic *S. symbiotica* may be more widespread across aphid genera in nature. The global distribution of *Aphis*-associated strains, along with their ability to transmit using the same transovarial route as *B. aphidicola*, suggests that related gut-associated strains may serve as a source pool for the evolution of commensal or mutualistic strains. Mutualism may have arisen several times independently in *S. symbiotica*, and, along with the subsequent horizontal transfer, would contribute to the phylogenetic discordance between facultative symbionts and their aphid hosts (54). The acquisition and replacement of secondary symbionts has occurred during the evolution of many ancient insect-microbe symbioses and may help hosts to escape the “evolutionary rabbit hole” of dependence on a primary symbiont that has become an ineffective mutualist due to genome decay (55, 56).

Transovarial transmission in aphids occurs by bacterial endocytosis into the syncytial cell of early embryos (8). What host and bacterial factors are involved in this pathway are unclear, but insights may be gained from other systems in which hosts are genetically tractable. In *Drosophila*, knockout of yolk proteins or the Yolkless receptor results in reduced localization to and/or endocytosis of *Spiroplasma* in embryos, suggesting that the vitellogenin pathway is involved in transovarial transmission (57). While parthenogenetic aphids do not undergo vitellogenesis or produce visible yolk, it is possible that similar receptor-mediated processes are used for *B. aphidicola* and *S. symbiotica* transmission, and that specific bacterial ligands are required. If this is the case, *S. symbiotica* strains that normally live in the aphid gut appear to possess the requisite molecular determinants, as they display an innate potential for endocytosis into embryos. Furthermore, the inability of *E. coli* BW25113 to endocytose into the syncytial cell of embryos suggests that the endocytic step of transovarial transmission contributes to selectivity in this system. While many bacterial taxa occasionally infect pea aphids (30, 58), few are found as long-term mutualists. The primary symbiont, *B. aphidicola*, is stably maintained in most aphid lineages, though rare replacements exist (e.g., (59)). Additionally, few species are found as secondary symbionts, and most are members of the *Enterobacterales*, including *S. symbiotica, Ca*. H. defensa, *Ca*. R. insecticola, *Ca*. F. symbiotica, and *Arsenophonus*, and, less commonly other bacterial groups, including *Wolbachia, Rickettsia*, and *Spiroplasma* (21, 22).

Aphids that are co-infected with *B. aphidicola* and mutualistic secondary symbionts possess several known mechanisms that limit the competition between these bacteria. For one, hosts can sort symbionts into distinct cell types, with *B. aphidicola* in primary bacteriocytes and *S. symbiotica* relegated to secondary bacteriocytes and sheath cells (60). Despite its pathogenicity, we observed that CWBI-2.3^T^ is not able to invade primary bacteriocytes with *B. aphidicola* and is compartmentalized into sheath cells in a similar manner as mutualistic strains (Figure 3). Pea aphid genotypes may vary in their ability to associate with secondary symbionts. In contrast to the results obtained with *Acyrthosiphon pisum* LSR1 in our study, when the facultative, host-restricted strain *S. symbiotica* IS was transferred to *Acyrthosiphon pisum* AIST, it showed a disordered localization, invading primary bacteriocytes with *B. aphidicola* (61). In these cases, *S. symbiotica* IS was trapped in primary bacteriocytes, unable to exocytose during transmission (8). The specific exocytosis of *B. aphidicola* also likely plays a role in limiting competition between *B. aphidicola* and secondary symbionts across multiple generations.

The vertical transmission of CWBI-2.3^T^ and HB1 in pea aphids is limited by their virulence in hemolymph, as infected aphids do not live long enough to birth offspring infected by transovarial transmission. Both CWBI-2.3^T^ and HB1 are more pathogenic in hemolymph than facultative *S. symbiotica* Tucson, but also notably far less pathogenic than *S. marcescens* Db11. Possibly, adaptation to the gut selects for reduced *Serratia* virulence by allowing *Serratia* the time to form a robust gut infection and transmit to other aphids, including offspring, via honeydew (40). The virulence of CWBI-2.3^T^ and HB1 coincides with unregulated titer, as both of these strains attain 100 to 1,000-fold higher titers relative to mutualistic *S. symbiotica* when injected into hemolymph. This enormous difference in titer, and the constancy of the low titers observed for the symbiotic strains, suggests that mutualistic *S. symbiotica* growth is regulated. The regulation of virulence and titer is important in the establishment of vertical transmission. Self-regulation of both titer and virulence through quorum sensing has been demonstrated in *Sodalis praecaptivus*, and allows this species to establish vertically transmitted infections in weevil and tsetse fly hosts (51, 62). Whether mutualistic *S. symbiotica* have relied on similar mechanisms to establish persistent vertical transmission in aphids is unclear. Alternatively, *S. symbiotica* virulence may be attenuated by the loss of one or several key virulence factors before the establishment of vertical transmission. The experimental tractability of these strains will allow for future investigations focused on these attenuation mechanisms and the role of vertical transmission in the transition to a host-restricted lifestyle in aphids.

## Materials and Methods

### Additional details are provided in SI Appendix

#### Isolation and culture of *S. symbiotica*

S. *symbiotica* strain CWBI-2.3^T^ (DSM 23270) was obtained from the DSMZ-German Collection of Microorganisms and Cell Cultures and was grown on tryptic soy agar (TSA) plates at 27 °C (25). *S. symbiotica* strain HB1 was isolated from the melon aphid (*Aphis gossypiĩ*), collected in August 2018 from HausBar Farms in Austin, Texas. Details are provided in SI Appendix.

#### *S. symbiotica* HB1 genome sequencing

*S. symbiotica* HB1 was grown in TSB at room temperature, harvested at an OD_600_ ~ 1.0, and DNA was extracted with the DNeasy Blood & Tissue Kit (Qiagen). A paired-end sequencing library with dual barcodes was prepared using the Illumina Nextera XT DNA kit, and sequencing was performed on an Illumina iSeq 100. Raw reads were trimmed using Trimmomatic (63) and assembled using the SPAdes algorithm (64) via Unicycler (65). Genome contamination and completeness were assessed using CheckM (66).

#### Phylogenetic analysis

*S. symbiotica* and outgroup genomes used for phylogenetic analysis are listed in Table S1. Genomes were downloaded from the NCBI Assembly Database on March 2, 2020. All outgroup genomes were filtered for completeness > 95% and contamination < 5% using CheckM (66). Annotations were obtained using Prokka (67), and 176 single-copy orthologs were identified by OrthoFinder (68). These single-copy orthologs were aligned with MAFFT (69), trimmed using a BLOSUM62 matrix in BMGE (70), and concatenated using an in-house script, producing an alignment with 56,881 total amino acid positions. A tree was constructed by maximum likelihood with a JTT+R10 model and 100 bootstraps, using IQ-Tree (71). The complete phylogeny is available as Figure S1. The presence of *Serratia* marker genes was determined using CheckM with the *Serratia* marker gene set provided with CheckM. The average nucleotide identity for *S. symbiotica* genomes was calculated using FastANI (72).

#### Chromosomal integration of sfGFP

Superfolder GFP (sfGFP) was integrated into the chromosomes of *S. symbiotica* CWBI-2.3^T^, *S. symbiotica* HB1, and *E. coli* BW25113 through mini-Tn7 tagging, as described by Choi *et al*. (73). Details are provided in SI Appendix.

#### Tracking aphid survival, fecundity, and transmission after injection with *S. marcescens, S. symbiotica*, and injection buffer

Fourth instar pea aphids were injected with *S. marcescens* Db11, recombinant *S. symbiotica* CWBI-2.3^T^-GFP, recombinant *S. symbiotica* HB1-GFP, hemolymph from pea aphids infected with *S. symbiotica* Tucson, or injection buffer, as described above. Every 24 h, survival was recorded, offspring were collected, and surviving adults were moved to a fresh dish. Adults were collected at death or at the end of the experiment at 15 DPI and screened for the presence or absence of *S. symbiotica*. Details are provided in SI Appendix.

#### Bacterial titer by spot-plating and quantitative PCR (qPCR)

Fourth instar pea aphids were injected with recombinant *S. symbiotica* CWBI-2.3^T^-GFP, recombinant *S. symbiotica* HB1-GFP, or with hemolymph from pea aphids infected with *S. symbiotica* Tucson, as described above. At 24 hours, aphids were transferred in sets of 15 to seedlings of *V. faba* and stored under long-day conditions (16H light, 8H dark) in incubators held at constant 20 °C. At each timepoint, aphids were collected in separate tubes, surface sterilized in 10% bleach for 1 minute, rinsed in deionized water for 1 min, then crushed and resuspended in 100 μL PBS. For aphids injected with culturable *S. symbiotica* CWBI-2.3^T^-GFP or HB1-GFP, 50 μL of this homogenate was used for spot plating and 50 μL frozen for DNA extraction and qPCR. For aphids injected with *S. symbiotica* Tucson, all 100 μL of homogenate was frozen and used for DNA extraction and qPCR. Details are provided in SI Appendix.

#### Statistical analyses

All statistical analyses and graphing were performed in the R programming language (version 3.6.3) (74). Survival rates for each treatment group were visualized as Kaplan-Meier survival curves, and comparisons of rates across treatment groups were performed using the Cox Proportional Hazards Model. Bacterial titers across treatment groups were compared using Kruskal-Wallis analysis of variance, followed by Dunn’s multiple comparisons test.

#### Fluorescence *in-situ* hybridization (FISH) microscopy

Fourth instar pea aphids were injected with wild-type *S. symbiotica* CWBI-2.3^T^, wild-type *S. symbiotica* HB1, or with hemolymph from pea aphids infected with *S. symbiotica* Tucson, as described above. Embryos were dissected at 4 DPI (Movie S1) or 7 DPI (Figure 3) in 70% ethanol. FISH was performed as in Koga *et al*. (8) with slight modifications. Details are provided in SI Appendix.

#### Live imaging of *E. coli* and *S. symbiotica* in pea aphids

For live imaging, fourth instar *Acyrthosiphon pisum* LSR1 were injected with recombinant *S. symbiotica* CWBI-2.3^T^-GFP or with recombinant *E. coli* BW25113-GFP, as described above. At 24 hours, aphids were transferred in sets of 15 to seedlings of *V. faba* and stored under long-day conditions (16H light, 8H dark) in incubators held at constant 20 °C. At 5 DPI, a subset of aphids from each treatment group were used to obtain titer counts via spot-plating, as described above, and remaining aphids were dissected in TC-100 insect medium. Single ovarioles from ten infected aphids per treatment group were observed under a Zeiss LSM 710 confocal microscope.

## Acknowledgements

We thank Kim Hammond for maintaining aphid lines, Dorsey Barger for access to insect collection sites at Hausbar Farms, and Margaret Steele, Travis Wiles, Peng Geng, and Sean Leonard for providing bacterial strains. We thank Ryuichi Koga for training and advice on FISH microscopy. We thank Daniel Deatherage for sequencing *S. symbiotica* HB1. We thank Anna Webb at the Microscopy and Imaging Facility of the Center for Biomedical Research Support at UT-Austin and Nicholas Kocian at Leica Microsystems for assistance with microscopy. We acknowledge the Texas Advanced Computing Center (TACC) for providing HPC resources. We thank members of Moran, Howard Ochman, and Barrick labs for helpful comments. pTNS2 was a gift from Herbert Schweizer (Addgene plasmids 64968). pTn7xKS-sfGFP was a gift from Karen Guillemin (Addgene plasmid 117394).

## Data availability

This Whole Genome Shotgun project for *S. symbiotica* HB1 has been deposited at DDBJ/ENA/GenBank under the accession JACBGK000000000. The version described in this paper is version JACBGK010000000.

## Competing interests

The authors declare that they have no competing interests.

## Funding

JP was supported in part by the University of Texas at Austin Provost Graduate Excellence Fellowship. This work was funded by the Defense Advanced Projects Agency (HR0011-17-2-0052 to JEB and NAM) and by the National Science Foundation (DEB-1551092 to NAM).

## Supplementary Figures, Tables, and Movies

**Figure S1.** Full phylogeny of *S. symbiotica* and outgroup. Maximum likelihood phylogeny with a JTT+R10 model and 100 bootstraps, based on concatenation of 176 single-copy orthologs (56,881 amino acid positions) shared across *S. symbiotica, Serratia, E. coli, Yersinia pestis*, and *Salmonella enterica*. Full list of genomes used for the phylogeny is provided in Table S1.

**Table S1.** Accession numbers, genome information, and references for analyzed *S. symbiotica* strains (Figures 1, S1).

**Table S2.** Average nucleotide identities for pairwise combinations of *S. symbiotica* strains.

**Movie S1.** Z-stack showing *S. symbiotica* HB1 entering the syncytium of a stage 7 embryo with *B. aphidicola* at 4 DPI. Blue, red, and green signal are used for host nuclei, *S. symbiotica*, and *B. aphidicola*, respectively.

**Movie S2.** Z-stack showing *S. symbiotica* CWBI-2.3^T^-GFP entering a live stage 7 embryo. Z-tack corresponds to the embryo depicted in Figure 4B.

**Movie S3.** Z-stack showing *S. symbiotica* CWBI-2.3^T^-GFP entering a live stage 7 embryo. Z-stack corresponds to the embryo depicted in Figure 4C.

**Movie S4.** Z-stack showing *E. coli* BW25113-GFP surrounding, but not entering, a live stage 7 embryo. Z-stack corresponds to the embryo depicted in Figure 4D.

**Movie S5.** Z-stack showing *E. coli* BW25113-GFP surrounding, but not entering, a live stage 7 embryo. Z-stack corresponds to the embryo depicted in Figure 4E.

## Notes

### Competing Interest Statement

The authors have declared no competing interest.

