## Supplementary material for "Vertical transmission at the pathogen-symbiont interface: *Serratia symbiotica* and aphids": SI Appendix

Supplemental Materials and Methods

**Isolation and culture of *S. symbiotica.*** S. *symbiotica* strain CWBI-2.3$^T$ (DSM 23270) was obtained from the DSMZ-German Collection of Microorganisms and Cell Cultures and was grown on tryptic soy agar (TSA) plates at 27 °C (1). *S. symbiotica* strain HB1 was isolated from the melon aphid (*Aphis gossypii*), collected in August 2018 from HausBar Farms in Austin, Texas. Individual aphids were surface-sterilized in 10% bleach for 1 min, rinsed in deionized water for 1 min, crushed with a sterile pestle in a 1.5 mL tube, resuspended in 100 µL PBS (137 mM NaCl, 10 mM Phosphate, 2.7 mM KCl, 8 mM $Na_2HPO_4$, 2 mM $KH_2PO_4$, pH = 7.4) and plated on TSA at 27 °C. After 10 days, small translucent colonies were visible and were individually transferred into tryptic soy broth (TSB). These liquid cultures were grown at room temperature with continuous shaking and reached an optical density at 600 nm ($OD_{600}$) of approximately 0.6 after 5 days. These were verified to be *S. symbiotica*, using 16S rRNA gene amplification and sequencing with primers 16SA1 and 16SB1 (2). Strains were cryopreserved in 80% glycerol and stored at −80 °C.

***S. symbiotica* HB1 genome sequencing.** *S. symbiotica* HB1 was grown in TSB at room temperature, harvested at an $OD_{600}$ ~ 1.0, and DNA extracted with the DNeasy Blood & Tissue Kit (Qiagen). A paired-end sequencing library with dual barcodes was prepared using the Illumina Nextera XT DNA kit and sequencing was performed on the Illumina iSeq 100. Raw reads were trimmed using Trimmomatic (3) and assembled using the SPAdes algorithm (4) via Unicycler (5). Genome contamination and completeness were assessed using CheckM (6).

**Phylogenetic analysis.** *S. symbiotica* and outgroup genomes used for phylogenetic analysis are listed in Table S1. Genomes were downloaded from the NCBI Assembly Database on March 2, 2020. All outgroup genomes were filtered for completeness > 95% and contamination < 5% using CheckM (6). Annotations were obtained using Prokka (7) and 176 single-copy orthologs were identified by OrthoFinder (8). These single-copy orthologs were aligned with MAFFT (9), trimmed using a BLOSUM62 matrix in BMGE (10), and concatenated using an in-house script, producing an alignment with 56,881 total amino acid positions. A tree was constructed by maximum likelihood with a JTT+R10 model and 100 bootstraps, using IQ-Tree (11). The complete phylogeny is available as Figure S1. The presence of *Serratia* marker genes was determined using CheckM with the *Serratia* marker gene set provided with CheckM. The average nucleotide identity for *S. symbiotica* genomes was calculated using FastANI (12).

**Chromosomal integration of sfGFP.** Superfolder GFP (sfGFP) was integrated into the chromosomes of *S. symbiotica* CWBI-2.3$^{T}$, *S. symbiotica* HB1, and *E. coli* BW25113 through mini-Tn7 tagging, as described by Choi *et al.* (13). We cloned sfGFP into a pUC18R6KT-miniTn7T-Gm backbone, provided by Wiles *et al.,* to produce pUC18R6KT-miniTn7T-Gm-sfGFP (14). *E. coli* MFDpir was used as a conjugal donor of pUC18R6KT-miniTn7T-Gm-sfGFP and of the helper plasmid pTNS2 (15). For triparental conjugation to recipient *S. symbiotica,* cultures were grown to an $OD_{600}$ of 0.6 – 1.2 and were combined in a 1:1:100 ratio of MFDpir with pUC18R6KT-miniTn7T-Gm-sfGFP, MFDpir with pTNS2, and *S. symbiotica,* respectively. Conjugation mixtures were plated in a 100 µL pool on TSA supplemented with 0.3 mM 2,6-diaminopimelic acid (DAP). After 24 hours, these conjugation mixtures were collected in 1 mL PBS, washed 3X in 1 mL PBS, and plated on TSA supplemented with gentamicin (10 µg/mL) and 1 mM Isopropyl β-D-1-thiogalactopyranoside (IPTG). Fluorescent colonies were observed after 5 days and were picked into liquid TSB supplemented with gentamicin (10 μg/mL), cryopreserved in 80% glycerol, and stored at -80 C. Triparental conjugation to recipient *E. coli* BW25113 was similar, but used a 1:1:1 ratio for 3 hours on LB + 0.3 mM DAP at 38 °C, outgrowth in LB, and selection on LB supplemented with gentamicin (10 μg/mL) and 1 mM IPTG. Transposon insertions were verified by PCR and sequencing with universal primer pTn7R and *Serratia*-specific PglmS-down-Ssymb (5'-GCACGTTGAAGAAATCGTAGC-3') or with pTn7R and *E. coli*-specific PglmS-down (13).

**Aphid rearing.** Aphids used in this study were cultivated as clonal isofemale lines reared on seedlings of broad bean (*Vicia faba*) under long-day conditions (16H light, 8H dark) in incubators held at constant 20 °C. Fourth instar *Acyrthosiphon pisum* LSR1 aphids were used as recipients for all injections. LSR1 was cured of *R. insecticola* in 2009 (16). Fourth instar *Acyrthosiphon pisum* Tucson were used as donors of *S. symbiotica* Tucson, a natural strain of *S. symbiotica* that has been maintained in this line for 21 years in the lab.

**Injection of *S. symbiotica*, *S. marcescens,* or *E. coli* to pea aphid hemolymph.** For injections of cultured *S. symbiotica* CWBI-2.3$^T$ or HB1, cells were grown in TSB at 27 °C with continuous shaking to an $OD_{600}$ of 0.5–1.0. For injections of *S. marcescens* Db11 or *E. coli* BW25113, overnight cultures were resuspended 1:100 in LB and grown with continuous shaking for 3 to 4 h at 37 °C to an $OD_{600}$ of 0.5–1.0. Prior to injection, cells were washed 3× in PBS, normalized to an $OD_{600}$ of 1.0, and diluted 100-fold in Buffer A (25 mM KCl, 10 mM $MgCl_2$, 250 mM sucrose, 35 mM Tris-HCl, pH 7.5). Approximately 0.1 μL of this resuspension was injected at the base of the hindleg, using beveled VWR 5 μL calibrated pipettes loaded onto a microinjector (Narishige IM-400) (MPa = 0.037, 0.2 seconds). To inject *S. symbiotica* Tucson, which is nonculturable and resides exclusively in pea aphid hemolymph, fourth instar *Acyrthosiphon pisum* Tucson aphids were surface sterilized in 10% bleach for 1 min, rinsed twice in deionized water for 1 min, crushed, and resuspended in 30 μL Buffer A. Approximately 0.1 μL of this resuspension was injected into recipients at the base of the hindleg. Injected aphids of the same treatment group were placed together on leaves of *V. faba* in Petri dishes to recover overnight and surviving aphids were used for subsequent data collection.

**Tracking aphid survival, fecundity, and transmission after injection with *S. marcescens, S. symbiotica,* and injection buffer.** Fourth instar pea aphids were injected with *S. marcescens* Db11*,* recombinant *S. symbiotica* CWBI-2.3$^T$-GFP, recombinant *S. symbiotica* HB1-GFP, hemolymph from pea aphids infected with *S. symbiotica* Tucson, or injection buffer, as described above. After 24 h, aphids from CWBI-2.3$^T$-GFP and HB1-GFP treatment groups were screened under blue light for the presence of GFP. GFP+ aphids from these treatment groups, along with the aphids from other treatment groups, were transferred to individual petri dishes containing a single leaf of *V. faba* inserted into a 1.5% water agar plug. 30 aphids were used for the *S. marcescens* Db11 treatment group, 20 aphids for the *S. symbiotica* CWBI-2.3$^T$-GFP, HB1-GFP, and Tucson treatment groups, and 27 aphids for the injection buffer control group. Petri dishes were stored under long-day conditions (16H light, 8H dark) at a constant 20 °C. Every 24 h, survival was recorded, offspring were collected, and surviving adults were moved to a fresh dish.

Adults were collected at death or at the end of the experiment at 15 DPI. Adults and offspring collected from CWBI-2.3$^T$ and HB1 treatment groups were crushed in a sterile pestle tube and resuspended in 50 µL PBS. Spot plating of 10 µL of this solution on TSA supplemented with gentamicin (10 µg/mL) and nystatin (100 U/mL) was used to determine the presence or absence of *S. symbiotica*. The remaining 40 µL was frozen at −20 °C as a safeguard. Adults and offspring collected from the Tucson treatment group were stored at −20 °C until screening. DNA of adults and offspring from the Tucson treatment group was extracted following the protocol of Bender *et al.* (17) and tested for presence or absence of *S. symbiotica* by PCR with primers PASScmp (5'-GCAATGTCTTATTAACACAT-3') and 16SA1 (5'-AGAGTTTGATCMTGGCTCAG-3') (18). Adults without confirmed cases of CWBI-2.3$^T$, HB1, or Tucson were filtered out of the dataset before analysis.

**Bacterial titer by spot-plating and quantitative PCR (qPCR).** Fourth instar pea aphids were injected with recombinant *S. symbiotica* CWBI-2.3$^T$-GFP, recombinant *S. symbiotica* HB1-GFP, or with hemolymph from pea aphids infected with *S. symbiotica* Tucson, as described above. At 24 hours, aphids were transferred in sets of 15 to seedlings of *V. faba* and stored under long-day conditions (16H light, 8H dark) in incubators held at constant 20 °C. At each timepoint, aphids were collected in separate tubes, surface sterilized in 10% bleach for 1 minute, rinsed in deionized water for 1 min, then crushed and resuspended in 100 µL PBS. For aphids injected with culturable *S. symbiotica* CWBI-2.3$^T$-GFP or HB1-GFP, 50 µL of this homogenate was used for spot plating and 50 µL frozen for DNA extraction and qPCR. For spot-plating, 10-fold dilutions from 1:10 to 1:10$^9$ were prepared in PBS in a 96-well plate. Spots of 10 µL were plated on TSA supplemented with gentamicin (10 µg/mL) and nystatin (100 U/mL). Fluorescent colonies were observed in 5-7 days and counted. For aphids injected with *S. symbiotica* Tucson, all 100 µL of homogenate was frozen and used for DNA extraction and qPCR. DNA extractions were performed with the DNeasy Blood and Tissue Kit (Qiagen), and qPCR reactions were performed in triplicate using iTaq$^{TM}$ Universal SYBR$^{®}$ Green Supermix (Bio-Rad) on an Eppendorf MasterCycler Realplex machine, with the following primers:

*Acyrthosiphon pisum ef1a* (XP_008182369) – 109 bp

Forward: 5'-gctgattgtgccgtgcttat-3' Reverse: 5'-cacccaaggtgaaagccaatag-3'

*S. symbiotica dnaK* (WP_006709606) – 124 bp

Forward: 5'-cttcacatcacccgccaatac-3' Reverse: 5'-gcctatggtgcagaagaaagtc-3'

**Statistical analyses.** All statistical analyses and graphing were performed in R version 3.6.3 statistical language (19). Survival rates for each treatment group were visualized as Kaplan-Meier survival curves, and comparisons of rates across treatment groups were performed using the Cox Proportional Hazards Model. Bacterial titers across treatment groups were compared using Kruskal-Wallis analysis of variance, followed by Dunn's multiple comparisons test.

**Fluorescence *in-situ* hybridization (FISH) microscopy.** Fourth instar pea aphids were injected with wild-type *S. symbiotica* CWBI-2.3$^{T}$, wild-type *S. symbiotica* HB1, or with hemolymph from pea aphids infected with *S. symbiotica* Tucson, as described above. Embryos were dissected at 4 DPI (Movie S1) or 7 DPI (Figure 3) in 70% ethanol. FISH was performed as in Koga *et al.* (20) with slight modifications. In brief, aphid ovaries were fixed overnight in Carnoy's solution (6:3:1 vol/vol ratio of ethanol:chloroform:acetic acid). Samples were washed with absolute ethanol, with PBS supplemented with 0.2% Tween$^{®}$20 (PBT), and with hybridization buffer (20 mM Tris-HCl (pH 8.0), 0.9 M NaCl, 0.01% SDS, 30% (vol/vol) formamide). Samples were incubated overnight in hybridization buffer containing 100 nM Cy3-PASSisR targeting 16S rRNA of *S.*

*symbiotica*, 100 nM Cy5-ApisP2A targeting 16S rRNA of *B. aphidicola*, and 0.5 µM SYTOX Green (Molecular Probes). Samples were washed in PBT, mounted in SlowFade™ Diamond Antifade Mountant (Molecular Probes), and observed under a Zeiss LSM 710 confocal microscope (*S. symbiotica* Tucson and *S. symbiotica* CWBI-2.3$^T$, Figure 3) or under a Leica TCS SP8 STED 3X microscope (*S. symbiotica* HB1, Movie S1).

**Live imaging of *E. coli* and *S. symbiotica* in pea aphids.** For live imaging, fourth instar *Acyrthosiphon pisum* LSR1 were injected with recombinant *S. symbiotica* CWBI-2.3$^T$ or with recombinant *E. coli* BW25113-GFP, as described above. At 24 hours, aphids were transferred in sets of 15 to seedlings of *V. faba* and stored under long-day conditions (16H light, 8H dark) in incubators held at constant 20 °C. At 5 DPI, a subset of aphids from each treatment group were used to obtain titer counts via spot-plating, as described above, and remaining aphids were dissected in TC-100 insect medium and embryos observed under a Zeiss LSM 710 confocal microscope.
